# Growth of *Sphagnum riparium* is strongly rhythmic: Contribution of the seasonal, circalunar and third rhythmic components

**DOI:** 10.1101/415539

**Authors:** Victor L. Mironov, Aleksei Y. Kondratev, Anna V. Shkurko

**Affiliations:** Institute of Biology KarRC RAS, Petrozavodsk, Russia; NRU Higher School of Economics, St. Petersburg, Russia, Institute for Problems of Regional Economics RAS, St. Petersburg, Russia; Lomonosov Moscow State University, Moscow, Russia

**Keywords:** growing season, growth monitoring, method of geotropic curvatures, synchronization, zeitgeber, lunar synodic cycle, cryptochromes

## Abstract

Continuous high-resolution monitoring of Sphagnum growth can provide insights into the biological rhythms of moss growth. Moss *Sphagnum riparium* is a convenient model for growth monitoring. Application of the method of geotropic curvatures has enabled a three-year monitoring with two to five-day intervals. We measured the increment in ca. 85000 shoots and produced ca. 3500 growth rate estimates, making this study a champion in precision compared to previous efforts. The zeitgeber for seasonal growth rhythms is the temperature seasonal cycle (R^2^=0.21–0.52). When the temperature changes by 10*°*C, moss growth rate is modified by 0.10–0.17cm/day according to the linear model, and 1.47–2.06-fold in the exponential model. The zeitgeber for circalunar rhythms is the lunar synodic cycle (R^2^=0.14–0.26). The average amplitude of the fluctuations it induces in the growth rate is 0.0425– 0.0572cm/day, which is equivalent to the effect of a 3.43–4.53*°*C change in temperature. The third rhythm can be distinguished in periodograms. Its period ranges from 10 to 16 days, but we did not detect the zeitgeber.

In total, three rhythms explain 51–78% of the growth rate. We believe that the strong rhythmicity in Sphagnum growth is associated with shoot growth synchronization.

## 1 Introduction

Sphagnum mosses are wetland engineers at high latitudes. They cover ca. 43% of the mire surface, mostly concentrating on raised bogs and transitional mires (Rydin et al., 2006; Vitt, 2006). The genus Sphagnum is represented by the calcifugous species (Vicherová et al., 2015) generating wet, acidic, and ion-poor conditions in wetland habitats, contributing to the active formation of peat (Clymo & Hayward, 1982; Rochefort, 2000). Peat mosses clearly tend to form carpets where shoots freely exchange nutrient solutions and efficiently retain moisture. In addition to precipitation and groundwater, other important sources of their mineral nutrition are symbiotic prokaryotes (Berg et al., 2013; Kostka et al., 2016) and translocation from old tissues (Rydin & Clymo, 1989; Aldous, 2002).

Overall, Sphagnum-dominated mires have a high significance in biospheric scale, since they:

1. constitute ca. 150mln. hectares of boreal and subarctic wetland habitats harboring unique diversity (Rydin et al., 2006),
2. participate in the global carbon cycle through the active sequestration of atmospheric carbon into peat (Gorham, 1991),
3. are active producers of iron chelates, which are essential for marine ecosystems (Krachler et al., 2016),
4. effectively biofilter methane, reducing its emissions by up to 50% (Raghoebarsing et al., 2005; Parmentier et al., 2011; Putkinen et al., 2018),
5. are powerful N_2_ fixing agents – this mechanism supplies up to 85–96% of the peatland’s total N budget (Larmola et al., 2014; Vile et al., 2014).

Sphagnum mosses grow strictly orthotropically, through division and the following elongation of apical meristem cells. They simultaneously decay at the bottom. As a result, 50 to 78% of the accumulated biomass is degraded in the acrothelm (Malmer & Wallén, 1993), and the remaining organic matter is transformed into peat. Growth in Sphagnum is confined by positive ambient temperatures, but substantial shoot elongation happens outside the traditional growing season (Laine et al., 2011). The growing season lasts the longest in waterlogged habitats, which freeze up later than drier ones (Hulme & Blyth, 1982).

On the global scale, the mean annual temperature taken alone explains 24–31% of Sphagnum production (Moore, 1989; Gunnarsson, 2005), while taken together with precipitation and geographical factors the level is 43% (Gunnarsson, 2005). Apart from temperature, moisture supply is a limiting factor for the growth of Sphagnum. Its constituents are total precipitation, number of days with precipitation, air humidity, as well as the physiological availability of moisture to plants, which is described using special climatic indices (Backéus, 1988; Asada et al., 2003). Acute moisture shortage leads to drought, which terminates many physiological processes, including growth.

Contemporary studies tend to look at the environmental aspect of Sphagnum growth, often relying on annual or near-annual increments. A far rarer approach is seasonal growth monitoring, but even there the time interval between individual observations is hardly ever less than 1.5–3 weeks and the combined duration of the observations is rather short (Sonesson et al., 1980; Murray et al., 1993; Gerdol, 1996; Asada et al., 2003). Paradoxically, researchers have not yet focused on continuous high-resolution monitoring (at *<*5-days intervals), although it yields the most precise data on moss growth. Our study highlights the ability of such monitoring to provide a new insight into the biological rhythm dimension of moss growth.

Biological rhythms encompass scales from individual cells to whole ecosystems, and their usual zeitgebers are circadian, annual (seasonal), and lunar geophysical cycles (Koukkari & Sothern, 2006). In terrestrial plants, circadian rhythms have been studied the most closely (McClung, 2006). Among bryophytes, some aspects of circadian rhythmicity are known from the model system *Physcomitrella patens* (Hedw.) Bruch & Schimp. (Aoki et al., 2004; Ichikawa et al., 2004; Holm et al., 2010). In contrast, infradian rhythms are far less known in terrestrial plants, and remain practically unknown in bryophytes.

We recently described a first case of infradian rhythmicity in the growth of the peat moss *Sphagnum riparium* Ångstr. (Mironov & Kondratev, 2017). Analysis of 11-day smoothed patterns of the growth rate of this species reveals a 30-day (circatrigintan) rhythm, where the coefficient r of the correlation between primary values of the growth rate (seasonal trend not removed) and the percentage illumination of the moon was r=-0.16 (n=1365; p*<*0.01) to r=-0.26 (n=530; p*<*0.01). Moving further in this direction, we managed to identify more rhythms and estimate their contribution to the growth of this moss.

In contrast to our previous paper (Mironov & Kondratev, 2017), the aim of this study is to analyze growth rate patterns in *S. riparium* from the perspectives of all rhythmic components. Data gathered by the geotropic curvatures method of our own design during a three-year high-precision growth monitoring effort were used to this end (Mironov, 2015; Mironov et al., 2016). In all, three rhythmic components have been identified, which collectively explain 51–78% of the moss growth rate. Two of the rhythms are associated with external zeitgebers (temperature and lunar synodic cycle), and one appears to be free-running. We believe that the significant collective contribution of rhythmicity to growth is associated with shoot growth synchronization and maintenance of the Sphagnum cover integrity.

## 2 Materials and methods

### Study area

The study was carried out in southern Karelia (Russia), in the middle taiga subzone (Aleksandrova & Yurkovskaya, 1989). The landscape under study area is the Shuiskaya lowland, which used to be the bottom of Lake Onego at the beginning of the Holocene. The prevalent Quaternary deposits there are varved clay and till (Demidov, 2010). Climatic conditions are outlined in Table 1.

The range for the study was a 0.12ha, herb-Sphagnum minerotrophic mire site (61*°*51’14"N; 34*°*10’51"E; 50m a.s.l.). Peat deposits at the site were 30cm thick at the edges, and 80cm in the central part. The water conditions during the growing season were as follows: mire water level – +2 to −20cm, TDS – 14 to 54mg/l, pH – 4.6 to 6.8, Eh – +168 to +240mV. Sphagnum mosses formed a nearly 100% continuous, intact cover. Dominant among them was *Sphagnum riparium*, which covered *>*90% of the area. There was a minor presence of *S. squarrosum* Crome, *S. magellanicum* Brid., *S. centrale* J.M. Black, *S. fallax* (H.Klinggr.) H.Klinggr., *S. angustifolium* Michx. Vascular plants grew unevenly, forming a 5 to 30% cover. The prevalent species among them were *Salix phylicifolia* L., *Equisetum fluviatile* L., *Calamagrostis canescens* (Web.) Roth, *Comarum palustre* L., *Carex rostrata* Stokes, *Calla palustris* L., *Typha latifolia* L.

**Table 1.** Climatic parameters in the study area

|  | Annual | The growing season |
| --- | --- | --- |
| T, °C | 3.1 | 9.9 |
| PR, mm | 611 | 427 |
| ND<sub>PR</sub>, days | 255 | 132 |
| CL<sub>T</sub>, % | 76 | 73 |
| CL<sub>L</sub>, % | 42 | 36 |
| C/CL/O, days | 16/152/197 | 9/104/101 |
| RH, % | 80 | 76 |

The following notations are used: T – mean temperature, PR – precipitation, ND<sub>PR</sub> – number of days with precipitation, CL<sub>T</sub> – total cloud cover, CL<sub>L</sub> – lower level cloud cover, C/CL/O – number of clear/cloudy/overcast days, RH – relative air humidity. The growing season is conventionally defined here as the interval from April 1 to October 31. The climatic parameters were calculated for the Petrozavodsk weather station (WMO ID 22820) using open-source data from www.rp5.ru.

### Study object

The object under study was a peat moss *S. riparium*. The species is dioecious, with a boreal and subarctic range (Daniels & Eddy, 1990). Communities dominated by this species are rather shortlived, wherefore the peat layers they generate are hardly ever thicker than 50cm (Galka et al., 2018). *S. riparium* is a common species in mire types that are transitional between rich fen and oligotrophic bog. It prefers wet habitats, which often appear as a result of secondary inundation. The prokaryotic N_2_-fixation associated with these conditions is at record-high values (Leppänen et al., 2015), and ca. 35% of the fixed N_2_ is supplied to Sphagnum shoots, intensifying its production (Berg et al., 2013). This is, probably, partly why *S. riparium* in wet habitats can have an annual increment of up to 42cm (Mironov et al., 2016), and productivity up to 1800g/m^2^ (Grabovik & Kuznetsov, 2016). Due to such passive protection against drought and high growth rates, this species appears to be a convenient model system for monitoring growth.

### Growth measurement

In monitoring surveys, where growth is measured over a short time interval, it is important that the measurement method does not affect the output. This requirement is best addressed by the application of innate biological markers of growth. Compared to artificial markers, they show a clear tendency for higher increment estimates, since deployment of the former may disturb the Sphagnum cover and inhibit shoot growth. According to Pouliot et al. (2010), increment demonstrated by innate markers would be 33–112% higher than when artificial markers are used. According to Mironov et al. (2016), this difference for *S. riparium* is 10–80%.

Here, we measured growth using geotropic curvatures, a method designed by the authors (Mironov, 2015; Mironov et al., 2016), and improved upon compared to its nearest analogs (Korchagin, 1960; Camill et al., 2001; Vitt, 2007). These innate markers are induced by a negative geotropic response to capitulum declination from the vertical. *S. riparium* shoots develop distinct geotropic curvatures, two varieties of which are used here – nival and impact-generated. The principle of their application in growth monitoring is shown in Fig. 1.

**Fig. 1.**
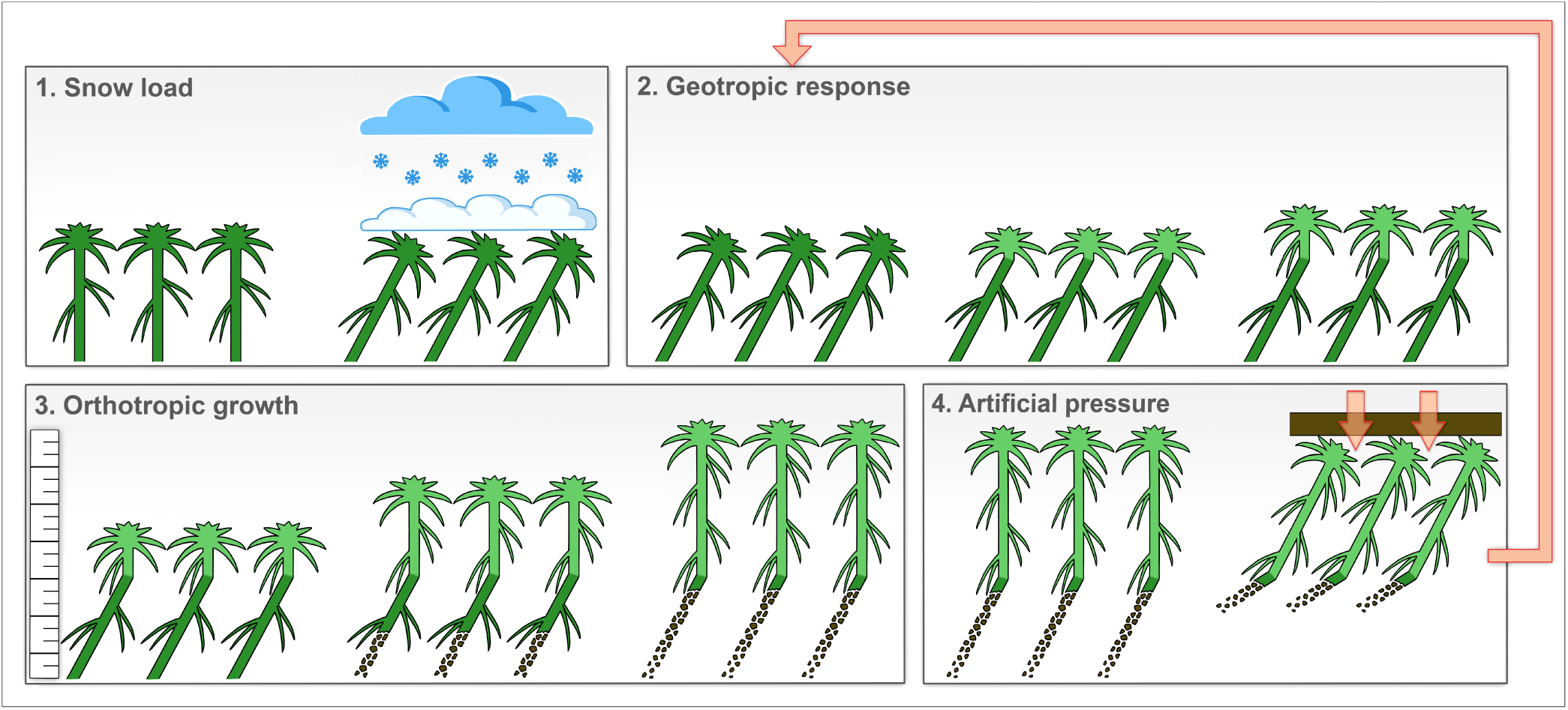
The principle of using geotropic curvatures for monitoring growth in *S. riparium*. 1. The snow cover compresses the Sphagnum carpet, making shoots deviate from the vertical.
2. When the temperature rises above zero, the geotropic response takes place, returning moss capitula to the vertical direction of growth. As a result, an innate growth marker is formed – the nival geotropic curvature – on the shoot.
3. After the position of the capitula has been rectified, shoots continue to grow, and the distance from the geotropic curvature to the apical meristem gradually increases. At this stage, growth is monitored. Monitoring the duration is limited by the decay of shoots at the bottom, which happens simultaneously with growth: if decay reaches up to the geotropic curvature, further monitoring becomes pointless.
4. To prevent curvature loss, we induce geotropic curvature in advance by artificially compressing the Sphagnum cover. This impact is analogous to the snow load effect. It re-starts the geotropic response and launches the formation of new markers on the stem – impact-induced geotropic curvatures (2) to enable the continuation of growth monitoring (3).

Originally, nival curvatures were used as markers. They appear in response to snow load, and are therefore widespread in boreal mires. From mid-summer, however, nival curvatures were observed to decay in our sample plots and could thus no longer be used as markers. In this case, we re-induced curvature formation. To this end, a 1m^2^ sheet of veneer was carefully placed on top of the Sphagnum carpet to create pressure. As a result, in 1–2 days the shoots developed impact-generated curvatures which we could then use as markers to further monitor growth. Thus, using a combination of nival and impact-generated curvatures, we managed to accomplish a continuous growth monitoring covering the entire growing season.

### Experimental design

The main parameters of the study are summarized in Tab. 2. Growth monitoring was carried out in a series of ca. 9m^2^ sample plots outlined within an *S. riparium* carpet with a minor presence of vascular plants (*<*15%). Every year, undisturbed portions of the carpet were chosen as sample plots. Monitoring started once the carpet had thawed (Apr. 11 in 2015, Apr. 14 in 2016, and Apr. 24 in 2017), and ended when it froze again (Oct. 5 in 2015, Oct. 12 in 2016, Oct. 20 in 2017). Thus, the survey embraced the entire period of *S. riparium* growth.

**Table 2.** Data volume in the study

| Year | 2015 | 2016 | 2017 |
| --- | --- | --- | --- |
| SD, days | 178 | 182 | 180 |
| NO, days | 34 | 68 | 88 |
| NSP | 4 | 11 | 13 |
| NSM | 9087 | 30267 | 45278 |
| NGREo | 95 | 505 | 789 |
| NGREe | 530 | 1365 | 1578 |
| MSS, shoots | 93.7 | 59.7 | 57.8 |
| MI, days | 5.2 ± 1.5 | 2.8 ± 0.8 | 2.0 ± 0.0 |

The following notations are used: SD – duration of the experiment, NO – number of observations, NSP – number of sample plots, NSM – number of shoots measured, NGREo – number of growth rate estimates, observed, NGREe – number of growth rate estimates, extrapolated, MSS – mean sample size, MI – mean interval between observations ± s.d.

Growth monitoring consisted of sequentially sampling shoots and calculating the increment difference from previously sampled shoots (Fig. 2). Shoots for increment estimation were taken from compact (ca. 100cm^2^) patches on the carpet. Each subsequent sampling was slightly (5–15cm) spaced off from the previous patch. Thus, for each sample plot we obtained, step-by-step, the pattern (time series) of the shoot increment, from which the growth rate was calculated. The growth rate is defined h ere as the p er d ay growth of shoots calculated by the formula:

**Fig. 2.**
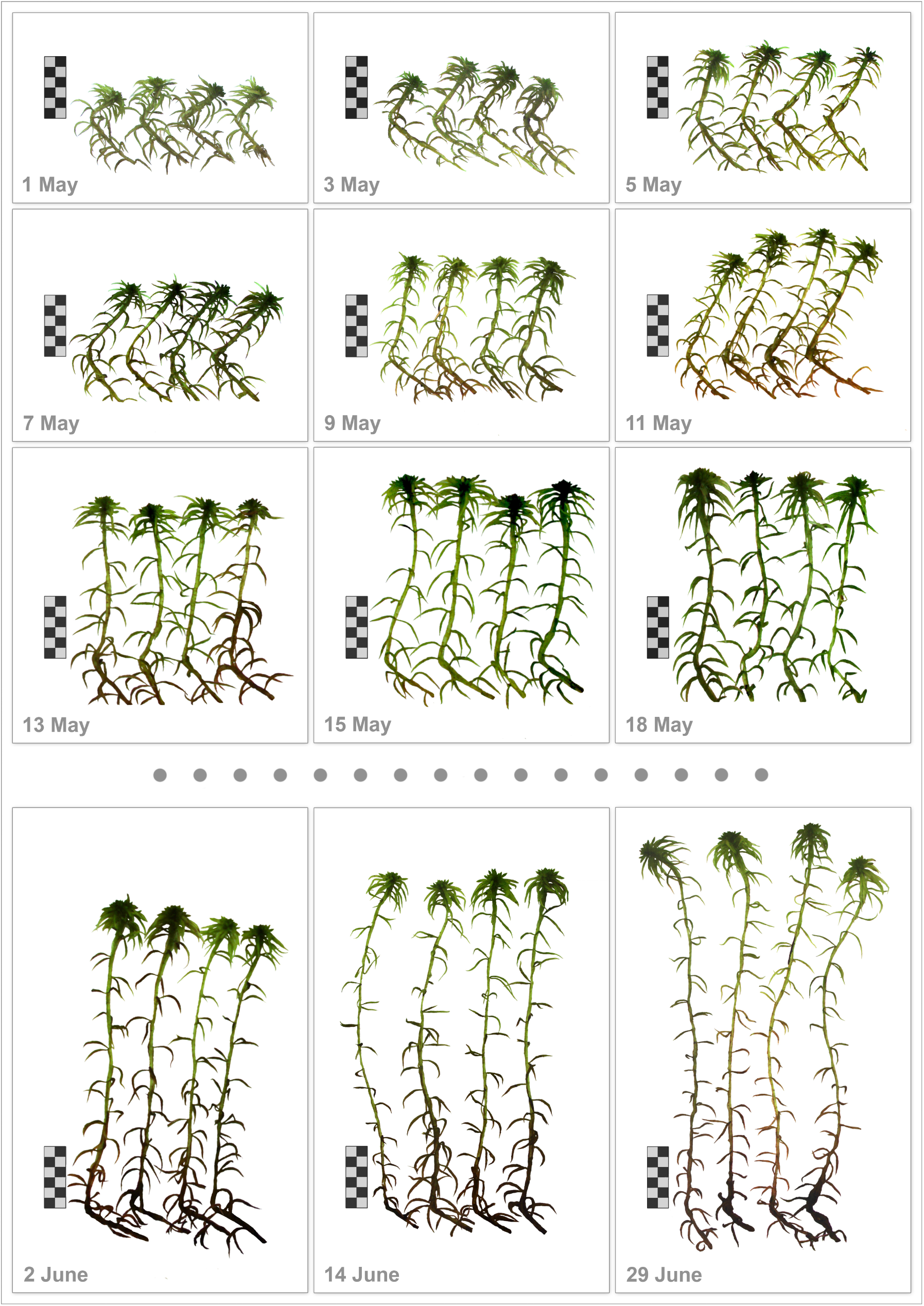
A fragment of *S. riparium* growth monitoring in one of the sample plots. Growth monitoring is shown for the period from May 1 to June 29, 2016. The length increment of shoots is the distance from the nival geotropic curvature (at the base of the scale marker) to the apical meristem. Scale interval is 5mm.

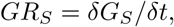

where *GR*_*S*_ is the growth rate in the sample plot, *δG*_*S*_ is the shoot increment, *δt* is the time interval. Based on *GR*_*S*_ values from all plots sampled within the same time interval, the mean growth rate across the experimental range *GR*_*α*_ was calculated by the formula:

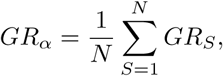

where *GR*_*α*_ is the mean growth rate in the experimental range,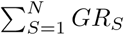 *GR*_*S*_ is the sum of growth rate values from all sample plots, *N* is the number of sample plots (corresponds to NSP from Tab. 2).

### Weather data and lunar synodic cycle profile

The study range is situated 4.5km away from the Petrozavodsk weather station (WMO ID 22820). A detailed archive of weather data from this station can be openly accessed online at www.rp5.ru. In this paper we use the mean daily temperature calculated using data from this weather station. To compute the exact profile of the lunar synodic cycle for the study area we used the Android freeware application Daff Moon Phase 2.87.

### Statistical data processing

This study employed standard statistical methods (P. Legendre & L.F. Legendre, 2012). The *GR*_*α*_ pattern for each year was transformed by moving the average smoothing with a three-day smoothing window. This procedure yielded a pattern of smoothed growth rate values *GR*_*β*_.

The *GR*_*β*_ pattern is used in the linear and exponential (van’t Hoff rule) models to estimate the temperature sensitivity of moss growth. The parameter of temperature sensitivity in the linear model is the coefficient K_1_, which demonstrates how much the growth rate is modified by a 1*°*C change in temperature. In the exponential model, temperature sensitivity is quantified by the coefficient Q_10_, which shows the fold change of the growth rate upon a 10*°*C change in temperature.

The pattern *GR*_*γ*_, which is the *GR*_*β*_ pattern after seasonal second-order polynomial trend removal, is used to investigate the effect of the lunar synodic cycle (LSC) on moss growth and to analyze rhythmic components. The time lag between moss growth and LSC is estimated by cross-correlation, and the extent of the LSC effect on moss growth is measured by linear regression. Rhythmic signals are studied by the spectral analysis (Lomb-Scargle periodogram). In doing so, we take into account not only the signal’s significance (P*<*0.01), but also the number of recurrences of each rhythm.

Both patterns *GR*_*β*_ & *GR*_*γ*_ are used to investigate the contribution of rhythmicity to moss growth (harmonic regression). For pattern *GR*_*β*_, we successively estimated the explanatory power of models with one (seasonal), two (seasonal + circalunar), and three (seasonal + circalunar + third), and for pattern *GR*_*γ*_ – with one (circalunar) and two (circalunar + third) rhythmic components. Data were statistically processed using PAST 3.19 and Excel 2007 software.

## 3 Results

### Growing season division

Fig. 3 shows the seasonal variation of the length increment in *S. riparium*, and Tab. 3 – the increment of shoots by months. The average growth rate (*±* s.d.) over the growing season was 0.23*±* 0.10cm/day in 2015, 0.21 *±*0.13cm/day in 2016, and 0.22*±* 0.12cm/day in 2017. The growing season falls quite distinctly into periods of latent and major growth (Tab. 4). According to our data, the duration of latent growth was 20–22 days, i.e., 12% of the growing season. We attribute it to the influx of meltwater from adjacent areas, which retards the warming up and development of the Sphagnum carpet. Once the meltwater is gone, shoots warm up quite quickly and the major growth period begins. In our study, its indicator is the steady transition to growth rates above 0.15cm/day. The major growth period lasted 157–162 days, i.e., 88% of the growing season.

**Fig. 3.**
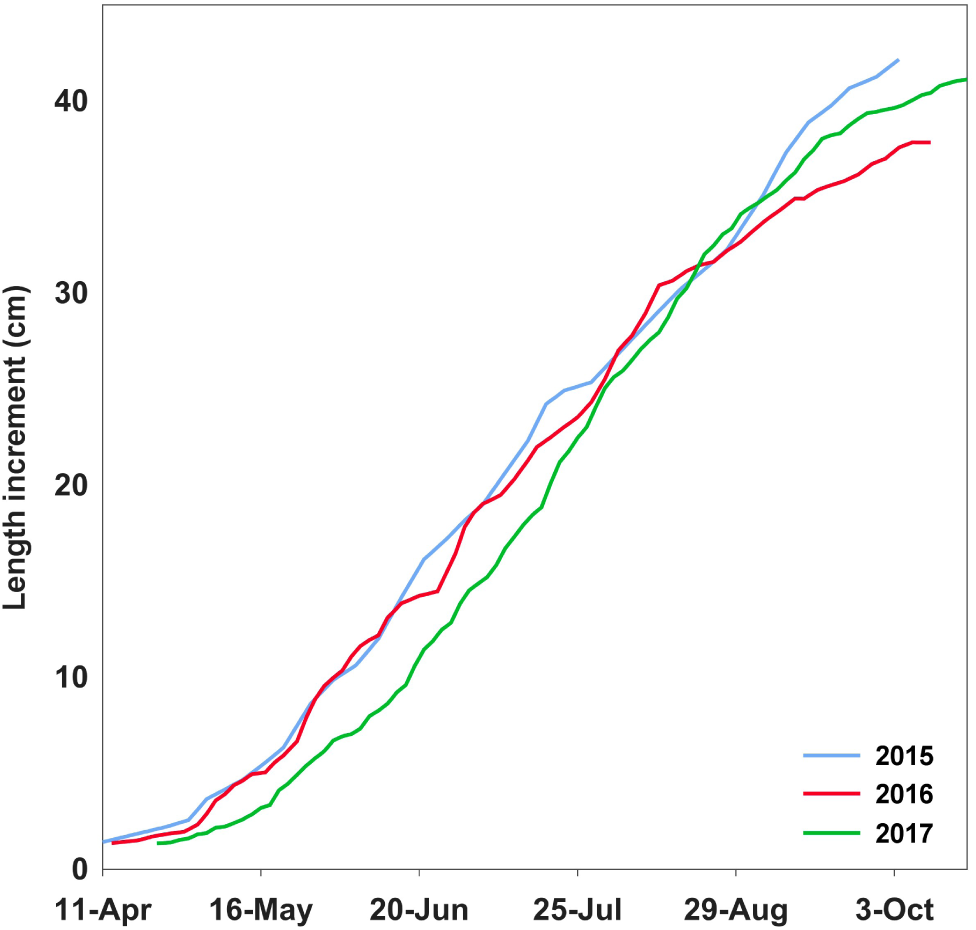
Cumulative growth of *S. riparium*. The curves were plotted using the growth rate patterns (*GR*_*β*_). The curves start from the increment values observed immediately after snowmelt – 1.43cm in 2015, 1.38cm in 2016, 1.36cm in 2017. The curves end at the values of increment over the entire growing season – 42.2cm in 2015, 37.9cm in 2016, 41.1cm in 2017.

**Table 3.**
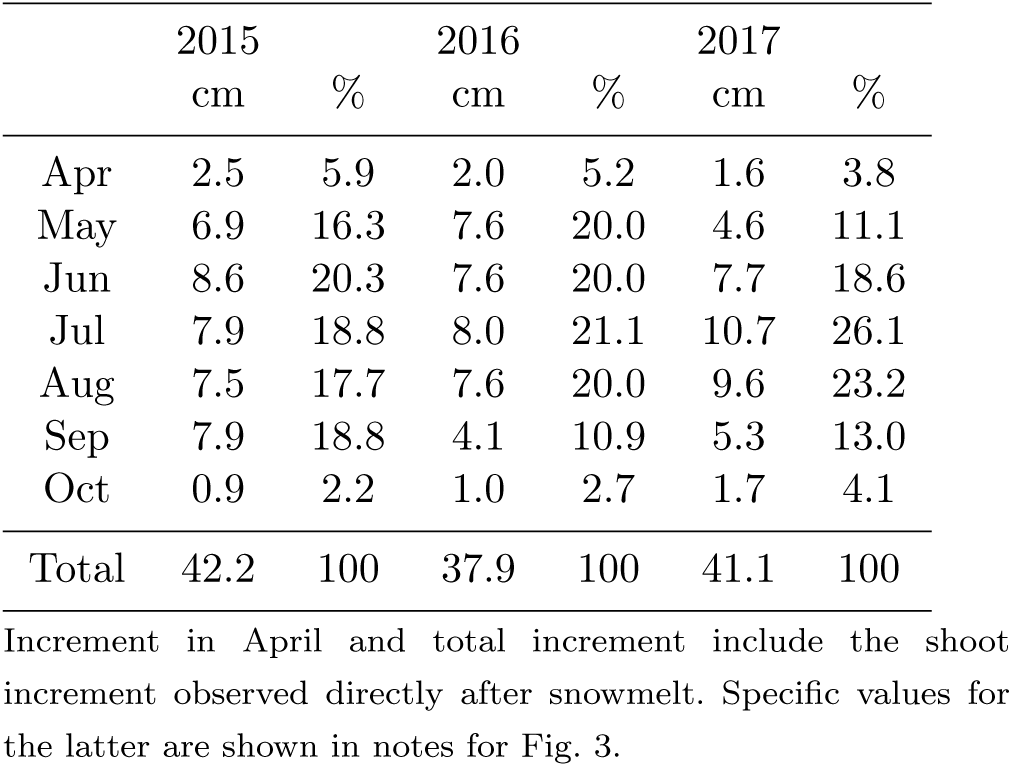
*S. riparium* growth distribution by months

**Table 4.** Characteristics of *S. riparium* growth periods

| Period,<br>Year | Interval | Duration,<br>days (%) | Growth rate,<br>cm/day±s.d. |
| --- | --- | --- | --- |
| latent | 11.04.–1.05. | 21 (11.8) | 0.05 ±0.02 |
| major | 2.05.–5.10. | 157 (88.2) | 0.25 ±0.10 |
| 2015 | 11.04.–5.10. | 178 (100) | 0.23 ±0.10 |
| latent | 14.04.–3.05. | 20 (11.1) | 0.05 ±0.04 |
| major | 4.05.–12.10. | 162 (88.9) | 0.22 ±0.14 |
| 2016 | 14.04.–12.10. | 182 (100) | 0.21 ±0.13 |
| latent | 24.04.–15.05. | 22 (12.2) | 0.07 ±0.04 |
| major | 16.05.–20.10. | 158 (87.8) | 0.24 ±0.13 |
| 2017 | 24.04.–20.10. | 180 (100) | 0.22 ±0.12 |

### Seasonal rhythmic component

The pattern *GR*_*β*_ matched the temperature seasonal cycle (Fig. 4). The time lag between temperature seasonal cycle trends and *GR*_*β*_ was one day in 2015, four days in 2016, and five days in 2017.

**Fig. 4.**
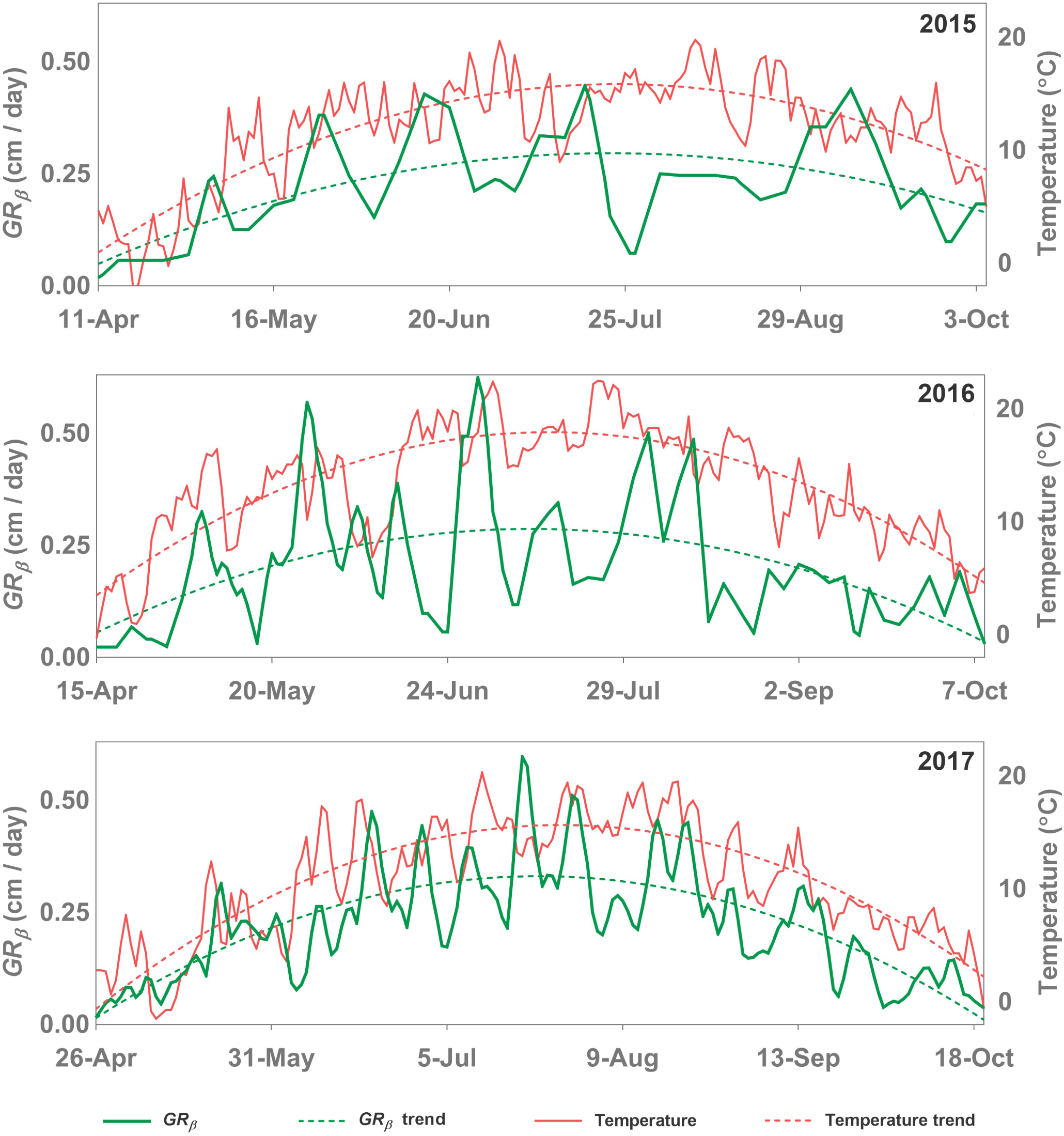
Seasonal variations of *S. riparium* growth rate (*GR*_*β*_) and mean daily air temperature. The trends in the figure are second-order polynomial trends plotted using the least squares method. The figure clearly demonstrates that the *GR*_*β*_ pattern follows the temperature seasonal cycle. Air temperature data come from the Petrozavodsk weather station.

The parameters of *GR*_*β*_ temperature sensitivity are summed up in Tab. 5. With a 10*°*C change in temperature, *GR*_*β*_ changed by 0.10–0.17cm/day in the linear model, and 1.47–2.06-fold in the exponential model. The linear model explained 21–52%, and the exponential model – 17–49% of *GR*_*β*_. Our data suggest that the modest explanatory power of the temperature models is due to the presence of non-seasonal rhythmic components in moss growth.

**Table 5.** Temperature sensitivity of *S. riparium* growth

| Linear | $K_1$ | $b$ | $n$ | $R^2$ |
| --- | --- | --- | --- | --- |
| 2015 | 0.010 | 0.11 | 178 | 0.21* |
| 2016 | 0.013 | 0.04 | 178 | 0.23* |
| 2017 | 0.017 | 0.05 | 178 | 0.52* |
| Total | 0.013 | 0.07 | 534 | 0.28* |
| Exponential | $Q_{10}$ | $GR_\beta(12)$ | $n$ | $R^2$ |
| 2015 | 1.47 | 0.23 | 178 | 0.17* |
| 2016 | 1.80 | 0.18 | 178 | 0.21* |
| 2017 | 2.06 | 0.23 | 178 | 0.49* |
| Total | 1.70 | 0.21 | 534 | 0.25* |

Linear model: $GR_\beta(t) = K_1 \cdot t + b$ , where $GR_\beta(t)$ is the growth rate (cm/day) at temperature $t$ , $K_1$ is the temperature coefficient indicating the change in growth rate at a 1°C change in temperature, $b$ is the free term. \* – $P < 0.01$ .
Exponential model: $GR_\beta(t) = GR_\beta(12) \cdot Q_{10}^{(t-12)/10}$ , where $GR_\beta(t)$ is the growth rate (cm/day) at temperature $t$ , $Q_{10}$ is the temperature coefficient indicating the fold change of the growth rate upon a 10°C change in temperature, $GR_\beta(12)$ is the growth rate at the temperature reference point set at 12°C. \* – $P < 0.01$ .

### Circalunar rhythmic component

*GR*_*γ*_ cross-correlation with LSC (Fig. 5a) reveals the presence of a significant negative correlation. We have r=-0.30 for 2015, r=-0.36 for 2016, and r=-0.43 for 2017, the time lag being zero. During the three years, the correlation peaked when *GR*_*γ*_ was slightly ahead of LSC. This time lead was three days (r=-0.38) in 2015, one day (r=-0.38) in 2016, three days (r=-0.52) in 2017. *GR*_*γ*_ showed a linear dependence on LSC (Fig. 5b), the explanatory power of the latter being 14–26%. The average LSC-induced amplitude of *GR*_*γ*_ was 0.0425cm/day in 2015, 0.0571cm/day in 2016, and 0.0572cm/day in 2017. The presence of a circalunar rhythmic component is verified by spectral analysis data (Fig. 6). The main peak of the signal in the periodograms corresponded to a 26-day rhythm in 2015, a 32-day rhythm in 2016, and a 29-day rhythm in 2017. The frequencies of these rhythms did not significantly differ from the LSC frequency.

**Fig. 5.**
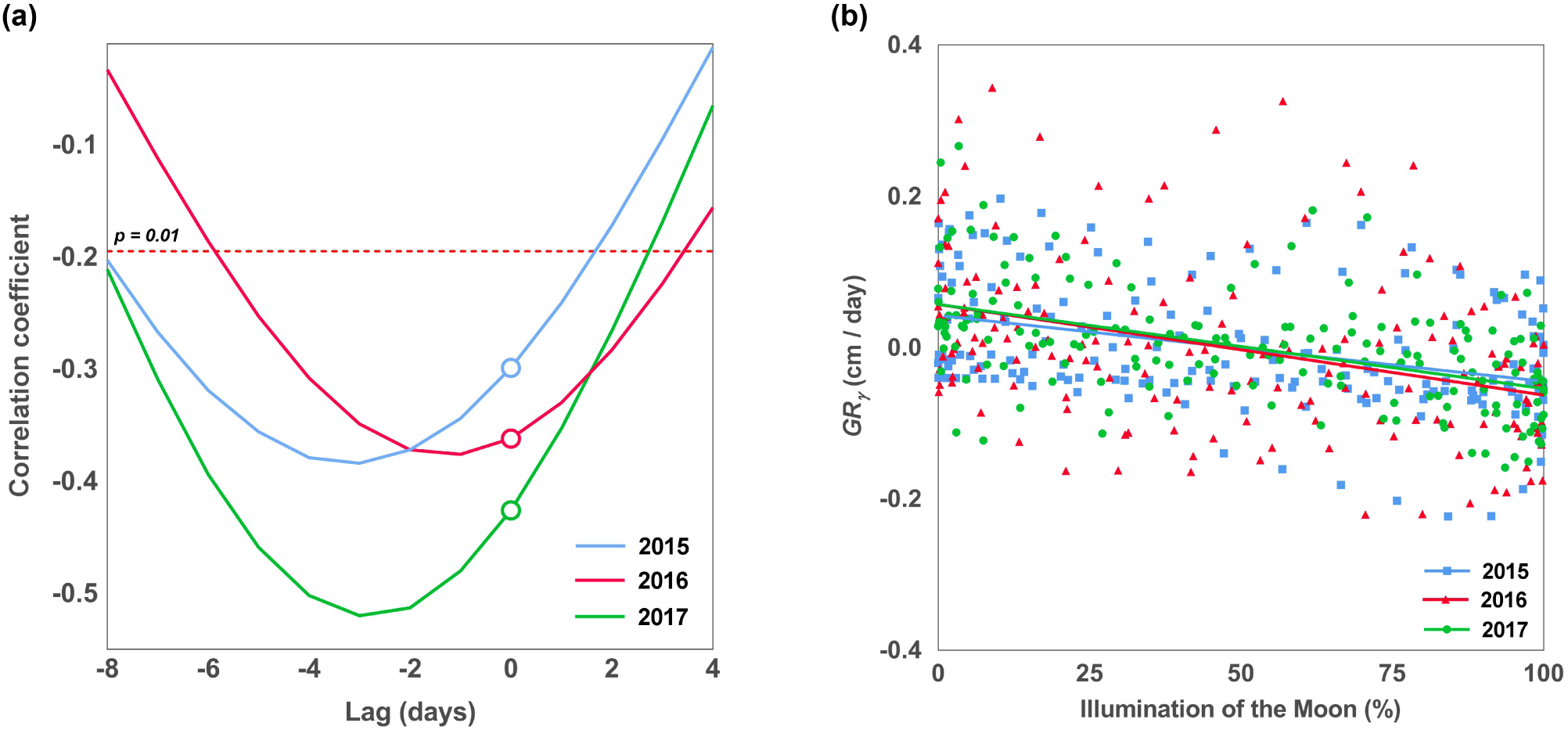
Cross-correlation (a) and linear regression (b) of detrended *S. riparium* growth rate (*GR*_*γ*_) and lunar synodic cycle. (a) Negative lag values correspond to a reciprocal shift of the time series, where *GR*_*γ*_ values are ahead of LSC values, and positive lag values – where *GR*_*γ*_ values lag behind LSC. Blank circles mark the values of the Pearson correlation coefficient at zero lag. Underneath the dash line is the region of significant (P*<*0.01) negative correlation. The size of the time series in the analysis was equalized to 178 data points for each of the three years. (b) The regression is for the lag of −2 days. The figure shows linear trends. For 2015 R^2^=0.14, for 2016 R^2^=0.14, for 2017 R^2^=0.26.

**Fig. 6.**
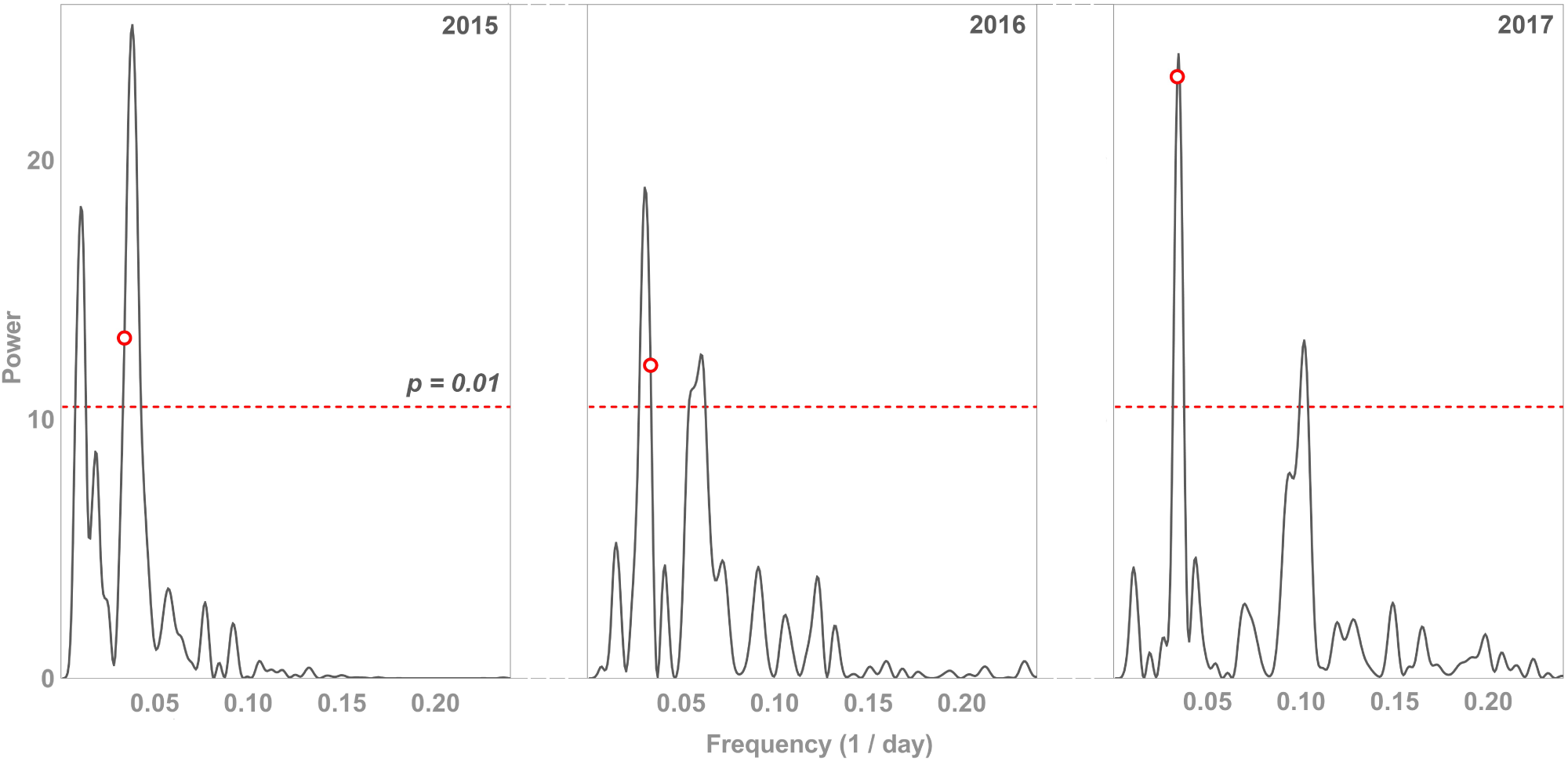
Lomb-Scargle periodograms of detrended *S. riparium* growth rate (*GR*_*γ*_) Above the dashed line is the region of significant (P*<*0.01) harmonic components. Significant frequencies correspond to 90-and 26-day rhythms in 2015, 32-and 16-day rhythms in 2016, 29-and 10-day rhythms in 2017. Blank circles mark the frequency corresponding to the lunar synodic cycle. It showed no significant differences from the frequencies of circalunar rhythmic components in any of the years.

### There is a third rhythmic component in *S. riparium* growth

In addition to the peaks corresponding to the circalunar rhythmic component, the periodograms contain other significant peaks.

The periodogram for 2015 features a peak corresponding to a 90-day rhythm. Having analyzed this peak, we found that this rhythm was caused by a fortnight-long slump of the growth rate in August, which concurred with a hot period with no rainfall and season’s lowest mire water level (−20cm). Hence, the rhythm could have been induced by water stress. This opinion is supported by the fact that this rhythm recurred only twice and was absent from the 2016 and 2017 periodograms. We therefore regard this 90-day rhythm as a pseudo-rhythm.

Periodograms for 2016 and 2017 contain significant peaks in the high-frequency region, which are absent from the 2015 periodogram because of a large monitoring interval. In 2016, the peak corresponded to a 16-day rhythm, and in 2017 – to a 10-day rhythm. Their corresponding rhythms recurred nine times in 2016 and sixteen times in 2017, and the analysis of their zeitgebers suggests neither of the rhythms was induced by temperature, geomagnetic field, or tidal activity.

### Rhythmicity contributes significantly to Sphagnum growth

The contribution of rhythmicity to Sphagnum growth was estimated by regressing the patterns *GR*_*β*_ and *GR*_*γ*_ with their harmonic models. The seasonal, circalunar and third rhythmic components were successively added to the *GR*_*β*_ pattern models (Fig. 7). The seasonal component alone explained 27–63% of *GR*_*β*_. The addition of the circalunar component raised the model’s explanatory power to 46–72%. Further addition of the third rhythmic component improved the explanatory power to 51–78%.

The contribution of rhythmicity excluding the dominant seasonal rhythm was estimated by using *GR*_*γ*_ pattern models to which the circalunar and the third rhythmic components were successively added (Fig. 8).

**Fig. 7.**
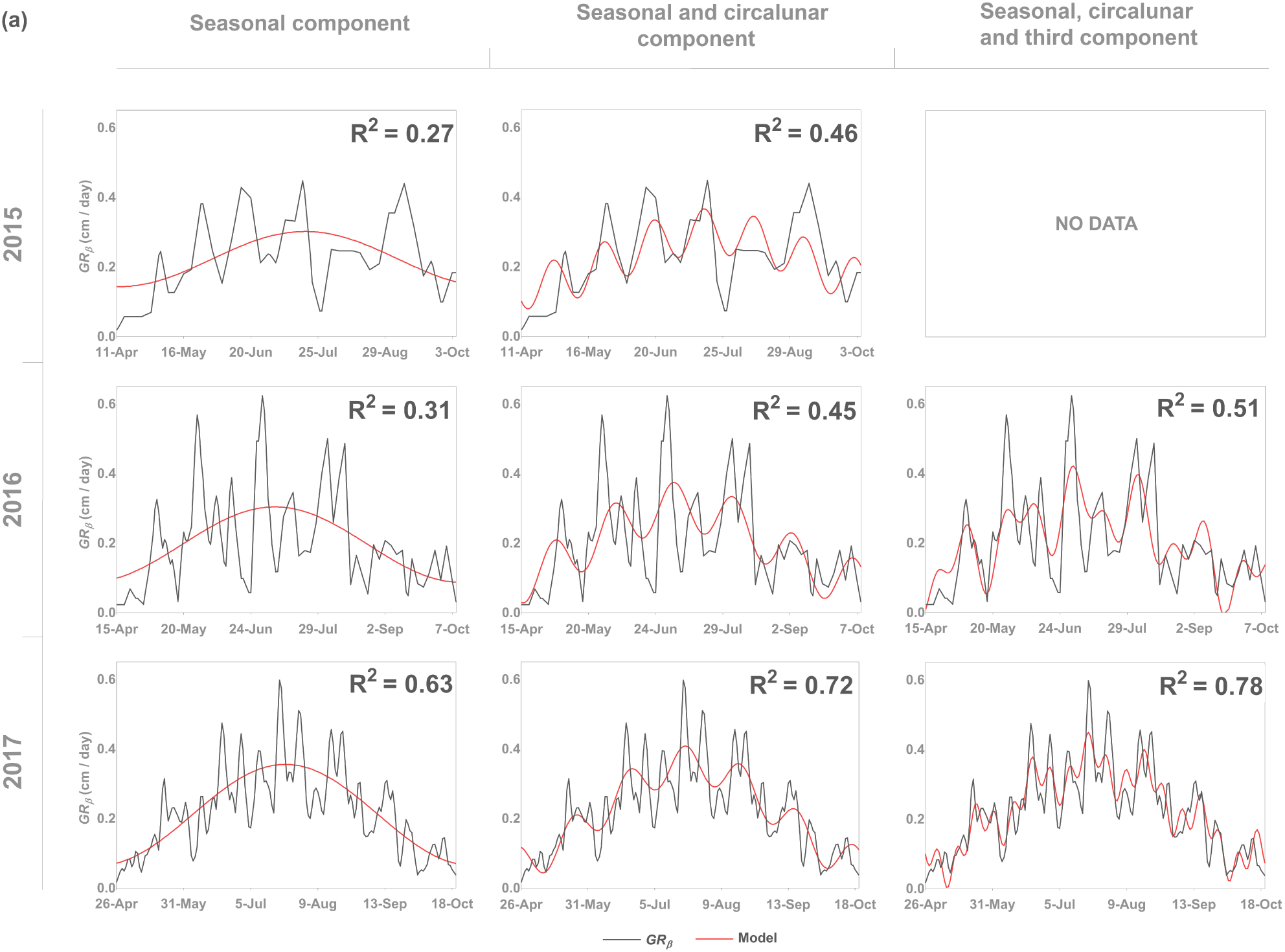
Harmonic models of the growth rate (*GR*_*β*_) for *S. riparium*.

**Fig. 8.**
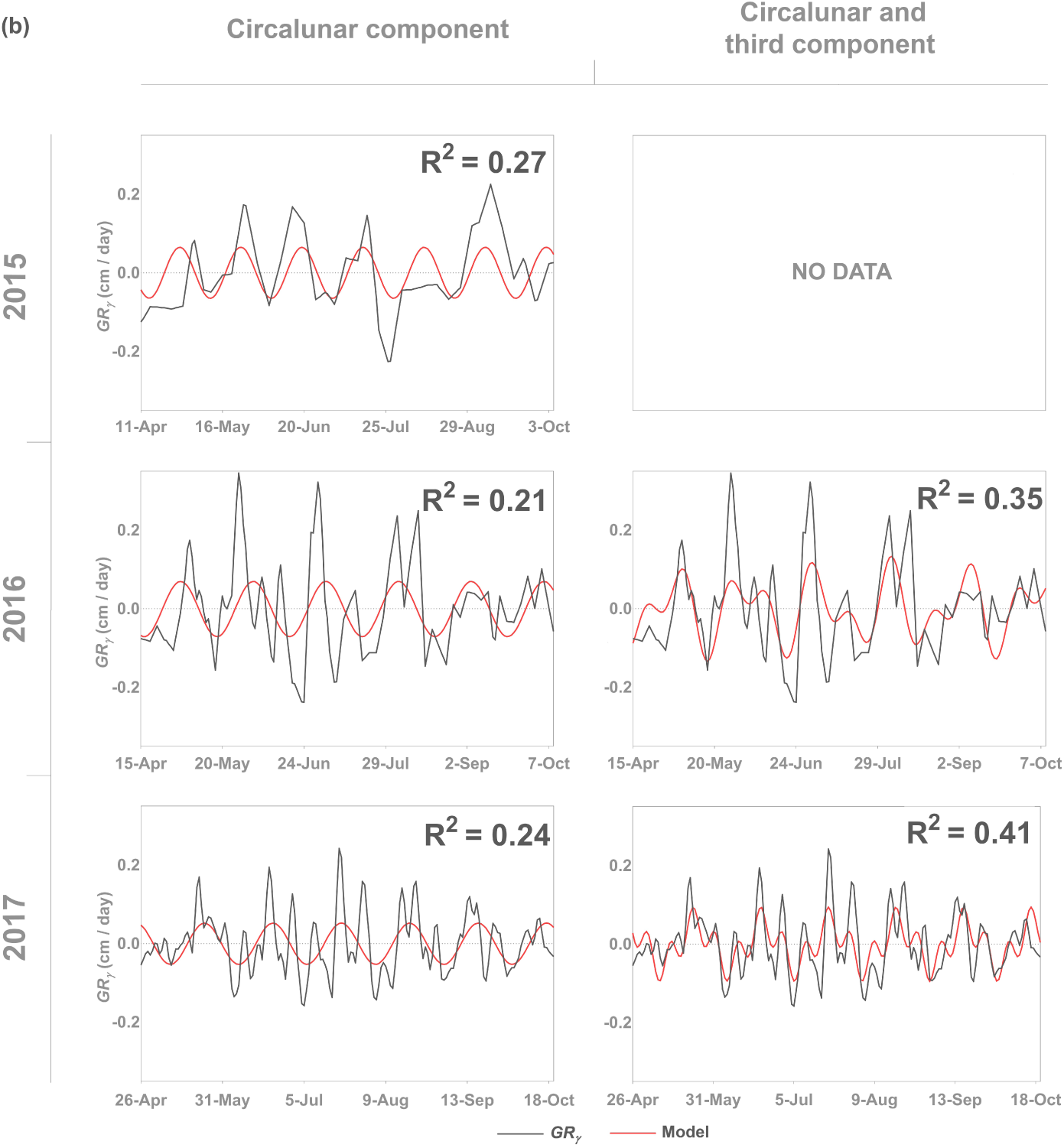
Harmonic models of the detrended growth rate (*GR*_*γ*_) for *S. riparium*.

The circalunar component alone explained 21–27% of *GR*_*γ*_, and the further addition of the third rhythmic component into the model increased its explanatory power to 35–41%. Thus, all three rhythms in the pattern *GR*_*β*_ collectively explained more than a half of the values, and two rhythms in the pattern *GR*_*γ*_ explained over one third of all values.

## 4 Discussion

The length increment of *S. riparium* in this study was 37.9 to 42.2cm, which is among record-high values for this species. A nearly identical increment was previously reported for *S. riparium* which was found growing 36km away from our range, in the drainage ditches of Rittusuo mire (Mironov et al., 2016). Similar increment levels are also known for other species of wet habitats (Overbeck & Happach, 1957; Brock & Bregman, 1989).

The growing season for *S. riparium* in this study was 178–182 days. At about the same latitudes in Finland, Lindholm (1990) found that the growing season for *S. fuscum* was 121–128 days, i.e., around 30% shorter than here. We attribute this tangible difference to the elongation of the growing season in wet habitats (Hulme & Blyth, 1982), and to a more comprehensive coverage owing to the method of geotropic curvatures that we are using. Furthermore, the growing season in our study may be protracted due to its vicinity close to an urban infrastructure (Zhang et al., 2004).

### Temperature dependence of seasonal rhythms

Seasonal patterns of plant development at high latitudes arise from the strong dependence of metabolism on temperature. In living systems, van’t Hoff’s rule works within the range of biologically relevant temperatures, and the coefficient Q_10_ for metabolic processes is usually within a range of 2 to 3 (Hegarty, 1973). According to our data, Q_10_ for *S. riparium* growth in the field is 1.47–2.06. This is slightly below the Q_10_ for Sphagnum photosynthesis under controlled conditions, which is 1.70–2.47 (Haraguchi & Yamada, 2011).

Van’t Hoff’s rule predicts the tendency for the formation of seasonal temperature-dependent patterns of physiological processes in bryophytes living at high latitudes. The seasonality of growth in bryophytes has been described by Longton & Greene (1969), Pitkin (1975), Hulme & Blyth (1982), Brock & Bregman (1989), and Rincon & Grime (1989), and the direct dependence of growth on temperature – by Furness & Grime (1982), and Moore (1989). Paradoxically, there are no previous reports of a clear direct correlation between the seasonal patterns of bryophyte growth and the seasonal temperature cycle. On the contrary, the opposite tendency for the seasonal temperature and growth rate has been detected for the case of epiphytic bryophytes (Pitkin, 1975).

Our study demonstrated a seasonal temperature-dependent growth rhythm of *S. riparium*. The most credulous explanation for the contradiction in the growth of *S. riparium* and epiphytic bryophytes (Pitkin, 1975) is probably the fundamental difference in moisture supply. Moisture supply to epiphytic bryophytes is poor and intermittent, and their growth is therefore mainly restrained by drought. Since the chance of drought is proportional to the temperature, one gets the impression that the seasonal temperature and growth rate follow opposite tendencies. In contrast to them, *S. riparium* grows in wet habitats and is therefore little affected by droughts. The growth-limiting factor in this case is the seasonal temperature cycle.

According to our results, temperature average contribution to *S. riparium* growth rate is 28%, which agrees with the previously estimated 24–31% contribution of the mean annual temperature to the annual increment in Sphagnum (Moore, 1989; Gunnarsson, 2005). This proves that the temperature input is maintained at the same level even where increment studies differ considerably in scale. On a large scale, as demonstrated by our data, the seasonal temperature cycle only sets the general trend for Sphagnum growth rate, whereas its local fluctuations deviate from the strict temperature dependence quite substantially. This happens because Sphagnum is exposed to multiple random factors in the field, and because there are non-seasonal rhythmic components in its growth.

### Relationship between circalunar rhythmicity and LSC

Terrestrial plants have demonstrated sensitivity to LSC in flowering and pollination (Rydin & Bolinder, 2015; Ben-Attia et al., 2016), electrical conductivity of tissue (Burr, 1947; Holzknecht & Zürcher, 2006), the content of moisture and elements in tissues (Vogt et al., 2002; Zürcher et al., 2010), shoot growth and movement (Maw, 1967; Abrami, 1972; Buda et al., 2001).

The data obtained in this study, together with previously published results (Mironov & Kondratev, 2017) are evidence of the sensitivity of *S. riparium* growth to LSC. It is noteworthy that this species is the first bryophyte found to follow circalunar rhythms in its growth. Like vascular plants (Maw, 1967; Abrami, 1972), *S. riparium* showed an acceleration of shoot growth around new moon, and a deceleration around full moon. We have estimated the average amplitude of such fluctuations to be equivalent to the effect of a 3.43–4.53*°*C change in temperature (Tab. 6).

**Table 6.**
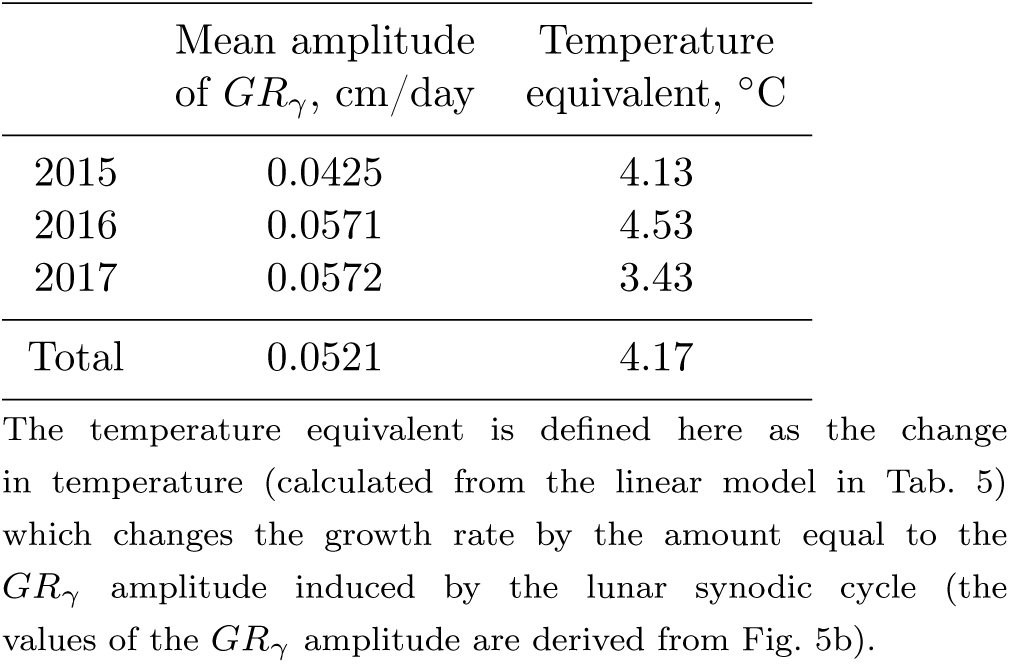
Temperature equivalent of the fluctuations of *S. riparium* detrended growth rate (*GR*_*γ*_) induced by the lunar synodic cycle

| | Mean amplitude<br>of $GR_\gamma$, cm/day | Temperature<br>equivalent, °C |
| --- | --- | --- |
| 2015 | 0.0425 | 4.13 |
| 2016 | 0.0571 | 4.53 |
| 2017 | 0.0572 | 3.43 |
| Total | 0.0521 | 4.17 |

To date, there are no definitive data on the mechanism of the lunar effect on plant growth, but the results of recent studies speak in favor of the cryptochrome-mediated mechanism of lunar signal reception. The main argument is the discovery made by Levy et al. (2007), that the cryptochrome gene expression differs significantly between full moon and new moon nights. Although the finding referred to the coral *Acropora millepora*, the high conservatism of cryptochrome genes gives us reason to believe that similar distinctions would occur among other plants, too.

The key interest here lies in the physiological reason for growth acceleration around the time of the new moon and its deceleration around full moon. Proceeding from the discovery of Imaizumi et al. (2002), the possible driver is the cryptochrome light signal-provoked suppression of auxin-induced gene expression, which results in growth inhibition. In our studies, such a negative growth response is associated with full moon periods, when lunar radiation and its cryptochrome sensitizing capacity are at a maximum.

At high latitudes, including our study area, the sensitivity of cryptochromes in plants to moonlight can be amplified by a geomagnetic field, which is stronger than at the Equator (Vanderstraeten et al., 2018). Furthermore, geomagnetic field fluctuations trigger cryptochro-me-mediated reactions in plants regardless of light stimuli (Agliassa et al., 2018). Given the obvious correlation of the geomagnetic field with LSC (Stolov & Cameron, 1964; Akimov & Dyatel, 2012), its modifications can act as an alternative zeitgeber where moonlight is poor or absent.

Thus, cryptochromes are at the same time sensitive to two signals that vary during LSC and are therefore the most suitable candidates for being used as moon biosensors.

### Does the third rhythmic component lack the zeitgeber?

In the patterns, we found only either a 16-day or a 10-day rhythm. It is therefore possible that they are modifications of the same rhythm. A likely suspect is a circadiseptan rhythm with a period of 11 to 17 days (Koukkari & Sothern, 2006), although the 10-day rhythm falls a little outside of this range. We are not aware of any previously made descriptions of such rhythms in bryophytes. Here, both rhythms are considered to be the third rhythmic component in Sphagnum growth. We have failed to identify its zeitgeber, but there is a chance of finding it in the future.

We have two hypotheses about why the period of the third rhythmic component was unsteady. Firstly, it could have been modulated by local environmental factors, such as weak magnetic fields (Martynyuk & Temur’ants, 2010) and thermal conditions (Kalita & Titlyanov, 2011). Secondly, the circalunar rhythm could have been the synchronizer for the third rhythmic component. A similar synchronization has been reported for beans, where circaseptan rhythmicity in the imbibition of dry stored bean seeds were found to be synchronized with the lunar synodic cycle (Spruyt et al., 1987). In our case, the third rhythmic component was a half of the circalunar cycle (16/32) in 2016, and one third of it (9.8/29.4) in 2017.

### Sphagnum carpet growth synchronization and high input from biorhythms

Sphagnum cover is an example of a supraorganism with a strong synchronization effect, which is needed to maintain the smooth surface of the carpet to minimize moisture losses. Biological rhythms are associated with the synchronization of functions at the cellular, organismal, and supra organismal levels, but the question remains of whether these rhythms are primary or secondary in relation to synchronization (Koukkari & Sothern, 2006).

Thus, the pronounced combined rhythmicity detected in our study (51–78% on patterns with the seasonal trend) is associated with *S. riparium* growth synchronization, but whether it is its cause or consequence remains debatable. The seasonal rhythmic component corresponds to the temperature-controlled synchronization of biological processes at the scale of the entire growing season (27–63% input to patterns with the seasonal trend). The circalunar and the third components are associated with growth synchronization at the scale of days and weeks (35–41% input to detrended patterns).

## 5 Conclusions

In this paper, Sphagnum growth is for the first time investigated from the rhythmicity perspective. This was made possible by implementing high-resolution growth monitoring based on a novel method of geotropic curvatures that we designed. We found the presence of the seasonal, circalunar and third rhythmic components in the growth of the moss.

The seasonal component was induced by the seasonal temperature cycle. The temperature sensitivity of growth in Sphagnum was estimated by the linear and the exponential (van’t Hoff rule) models: a 10*°*C change in temperature modifies the moss growth rate by 0.10–0.17cm/day in the linear model (R^2^=0.21–0.52) and 1.47–2.06-fold in the exponential model (R^2^=0.17– 0.49).

The circalunar component was induced by the lunar synodic cycle. As opposed to our previous study, where only an 11-day linear smoothing without seasonal trend removal was used (Mironov & Kondratev, 2017), here we offer an in-depth analysis of circalunar rhythmicity. After the seasonal trend was removed, the Sphagnum growth rate demonstrated a linear dependence on the illumination of the moon (R^2^=0.14–0.26), this correlation being the highest when LSC lagged slightly behind growth. The mean amplitude of moon-induced growth fluctuations was equivalent to a 3.43–4.53*°*C change in temperature. Relying on the results of ground-breaking studies of photoreceptors cryptochromes, we hypothesize that the mechanism of circalunar rhythmicity may be rooted in the sensitivity of cryptochromes to both moonlight and moon-induced variations in the geomagnetic field.

The third rhythmic component comes out in periodograms when the monitoring interval is reduced to less than three days. Its period is a multiple of the circalunar rhythm, and lasts 10–16 days. We found no zeitgeber for the third rhythm, but there may be some chance of spotting it in the future. This question remains open for further studies.

We have estimated in this study that the three rhythmic components collectively contribute 51–78% to Sphagnum growth, and two components (excluding the seasonal one) – 35–41%. The unexpectedly high input of rhythmicity to growth in bryophytes is reported for the first time here. This phenomenon is associated with the synchronization of Sphagnum carpet growth, but whether it is the cause or the consequence of this synchronization is still debatable.

We believe our results will encourage scientists to implement high-precision studies of growth in bryophytes, and design new methods for such studies. Since the rhythmic aspect of growth has not yet been investigated, we suggest it is taken into account in further research on growth processes among bryophytes.

